# Digital profiling of cancer transcriptomes from histology images with grouped vision attention

**DOI:** 10.1101/2023.09.28.560068

**Authors:** Yuanning Zheng, Marija Pizurica, Francisco Carrillo-Perez, Humaira Noor, Wei Yao, Christian Wohlfart, Kathleen Marchal, Antoaneta Vladimirova, Olivier Gevaert

**Author notes:** These authors contributed equally to this work (Yuanning Zheng, Marija Pizurica, Francisco Carrillo-Perez).

## Abstract

Cancer is a heterogeneous disease that demands precise molecular profiling for better understanding and management. Recently, deep learning has demonstrated potentials for cost-efficient prediction of molecular alterations from histology images. While transformer-based deep learning architectures have enabled significant progress in non-medical domains, their application to histology images remains limited due to small dataset sizes coupled with the explosion of trainable parameters. Here, we develop ***SEQUOIA***, a transformer model to predict cancer transcriptomes from whole-slide histology images. To enable the full potential of transformers, we first pre-train the model using data from 1,802 normal tissues. Then, we fine-tune and evaluate the model in 4,331 tumor samples across nine cancer types. The prediction performance is assessed at individual gene levels and pathway levels through Pearson correlation analysis and root mean square error. The generalization capacity is validated across two independent cohorts comprising 1,305 tumors. In predicting the expression levels of 25,749 genes, the highest performance is observed in cancers from breast, kidney and lung, where ***SEQUOIA*** accurately predicts the expression of 11,069, 10,086 and 8,759 genes, respectively. The accurately predicted genes are associated with the regulation of inflammatory response, cell cycles and metabolisms. While the model is trained at the tissue level, we showcase its potential in predicting spatial gene expression patterns using spatial transcriptomics datasets. Leveraging the prediction performance, we develop a digital gene expression signature that predicts the risk of recurrence in breast cancer. ***SEQUOIA*** deciphers clinically relevant gene expression patterns from histology images, opening avenues for improved cancer management and personalized therapies.

## Introduction

Cancer is a dynamic disease characterized by intricate molecular and cellular evolution. Over the course of evolution, cancer becomes more heterogeneous, and this heterogeneity is classified into inter-patient heterogeneity and intra-tumoral heterogeneity [1]. Inter-patient heterogeneity refers to the difference found between patients, which results from patient-specific factors, including germline genetic variations, differences in mutation profiles and environmental factors [2–4]. Comparatively, intra-tumoral heterogeneity describes the co-existence of cell subpopulations carrying different genomics, epigenomics and transcriptomics profiles within the same tissue. The spatial distribution of these cell subpopulations forms a complex ecosystem fostering signaling transduction that drives tumor progression [5, 6]. A systematic understanding of cancer heterogeneity presents formidable challenges for effective diagnosis and management.

With the growing interest in precision medicine, molecular profiling has gained significant attention as a critical component of prognostication and treatment planning. In the past decade, the advancement of RNA sequencing (RNA-seq) has enabled the comprehensive measurement of gene expression profiles at both bulk tissue levels and at locoregional levels [3, 5]. The resulting information has deepened our understanding of cancer heterogeneity, leading to the discovery of molecular signatures associated with treatment sensitivity [7–9]. However, incorporating gene expression analysis into clinical practice still represents a challenge. Current methods involve time-consuming and expensive laboratory procedures, limiting the integration of gene expression analysis in routine diagnostics.

With the digitization of histopathology glass slides into Whole Slide Images (WSIs), unprecedented opportunities arise for cost-efficient analyses of tumor properties. Notably, WSIs are available without additional cost as they are obtained in routine clinical practice for diagnostics. Despite providing only morphological information, WSIs can also reflect the molecular status of tumors. Over the past decade, deep learning-based computational methods have been developed to extract hidden morphological features from WSIs that associate with molecular properties. Convolutional neural networks (CNN) are trained to predict aneuploidies, genetic alterations and expression signatures of cancer infiltrating immune cells from WSIs [10–19].

Although remarkable progress has been made in computer vision for medical images, applying state-of-the-art methods to WSIs remains exceedingly challenging. Due to their immense size and resolutions, WSIs are first cropped into thousands of smaller tiles to enable feature extraction [12, 15, 16, 18, 20, 21]. The lack of fine-grained annotations for individual tiles poses significant challenges in extracting relevant morphological information associated with slide-level molecular features. Moreover, tile-level models cannot capture contextual and hierarchical relationships between multiple tiles of an image.

Recently, transformer-based deep-learning architectures have been adapted for computer vision [22]. Compared to traditional CNN models, vision transformers employ attention-based mechanisms to capture the contextual relationships between different regions of an image. Vision transformers have been trained on histology images for phenotype classification, disease subtyping and survival predictions [23, 24]. Alsaafin et al. developed tRNAsformer, a deep-learning method using attention-based mechanism to infer gene expression levels from WSIs [25]. This study provides evidence that transformers can learn high-dimensional molecular features from WSIs. However, the study is limited in three critical aspects. First, the model is trained exclusively on data from kidney cancer, and its applicability to cancers from other origins remains to be elucidated. Second, due to explosion in the number of trainable parameters, vision transformers have been shown to excel particularly when the model has been pretrained on large-scale datasets [26]. However, the benefit of pretraining was not integrated into their models. Third, the study only investigates the capacity of transformers in predicting gene expression levels at bulk tissue levels, whereas effective computational methods for resolving intra-tumoral gene expression heterogeneity are still limited.

Here, we present *SEQUOIA*, a deep learning model for **S**lide-based **E**xpression **Qu**antification using gr**o**uped V**i**sion **A**ttention. To generate contextualized representations of features from WSIs, we employ the transformer architecture to implement self-attention. The model is trained to automatically derive which tiles are relevant for the slide-level gene expression prediction. To enable the full potential of transformers, we pretrained the transformer encoder using 1,802 slides from six normal tissues. The model is then fine-tuned and evaluated using data from 4,331 tumor samples across nine cancer types. To assess the generalization capacity of our models, we validated our predictions in two independent cohorts. Through comprehensive biological pathway analysis, we demonstrate the model’s capacity to accurately predict gene expression governing inflammatory response, cancer cell metabolism and proliferation. Moreover, we develop a computational technique that utilizes our models to resolve spatial gene expression patterns within tumor tissues. Finally, by developing a novel RNA-seq-based gene expression signature, we demonstrate the clinical utility of our model in predicting breast cancer recurrence. *SEQUOIA* offers a cost-efficient way to infer and analyze gene expression patterns on a large scale, with potential applications in both research and clinical settings.

## Results

### *SEQUOIA* as tool for gene expression prediction from WSIs

We present *SEQUOIA*, a deep learning model for **S**lide-based **E**xpression **Qu**antification using gr**o**uped v**i**sion **A**ttention (“Methods”, Figures 1a-b-c). To train and evaluate the model, we utilized WSIs and RNA-seq gene expression data of nine cancer types available in The Cancer Genome Atlas (TCGA, Supplementary Table A1): (1) prostate adenocarcinoma (PRAD), (2) pancreatic adenocarcinoma (PAAD), (3) lung adenocarcinoma (LUAD), (4) lung squamous cell carcinoma (LUSC), (5) kidney renal papillary cell carcinoma (KIRP), (6) kidney renal clear cell carcinoma (KIRC), (7) glioblastoma multiforme (GBM), (8) colon adenocarcinoma (COAD), and (9) breast invasive carcinoma (BRCA).

**Fig. 1:**
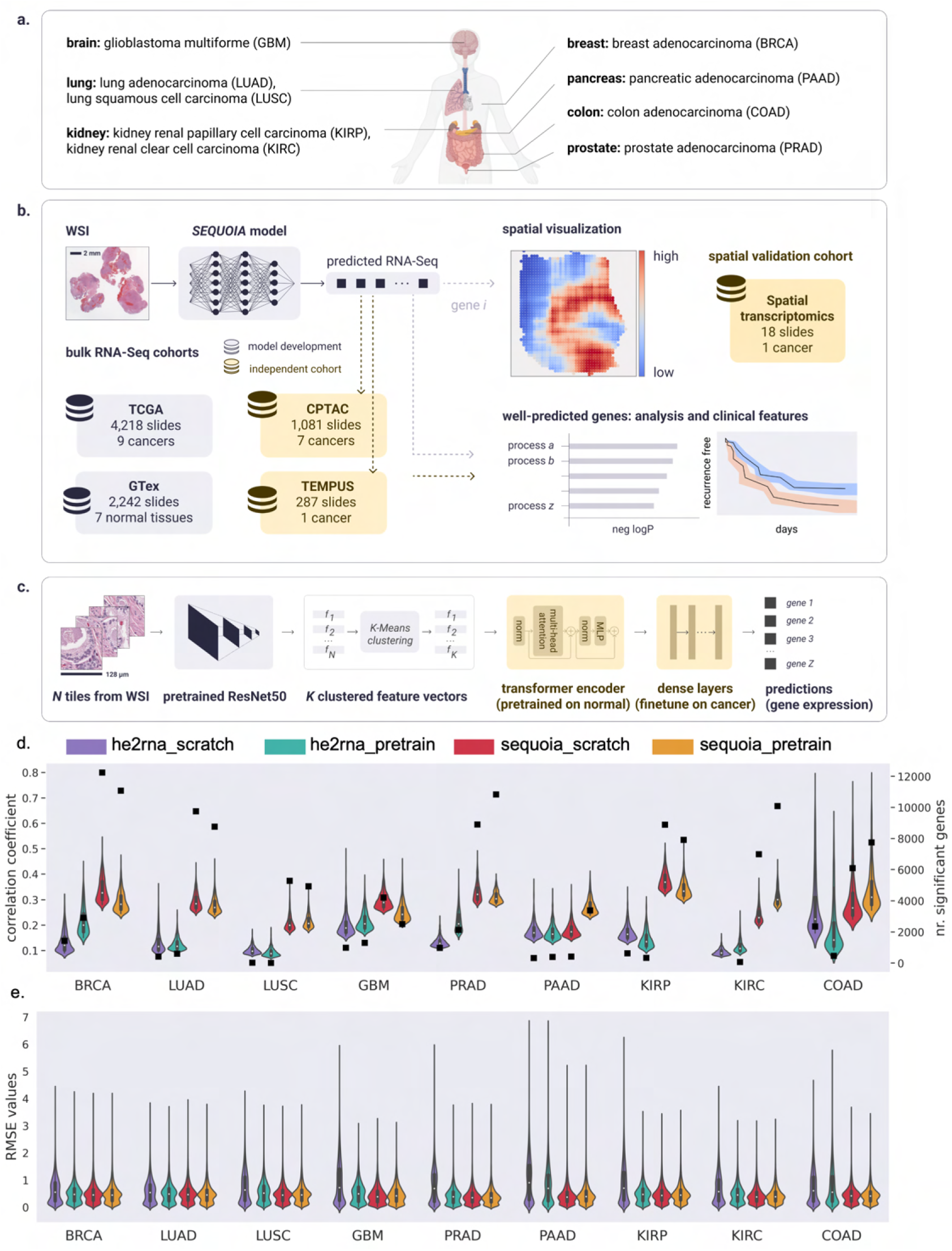
Overview of the workflow for the *SEQUOIA* model. a) Cancer types on which the *SEQUOIA* model is developed and validated. The panel is created with BioRender.com. b) The model is trained and evaluated using matched WSIs and bulk RNA-Seq data from nine cancer types available in the TCGA database. To pretrain the transformer encoder, we use matched WSIs and gene expression data of normal tissues from the GTEx database. The model is independently validated using data from the CPTAC and Tempus cohorts. Apart from predicting tissue-level gene expression, we integrate a spatial prediction technique that elucidates region-level gene expression patterns within tumor tissues, validated using a spatial transcriptomics dataset [5]. Clinical utility is demonstrated by evaluating the model’s capacity to predict cancer recurrence. c) *SEQUOIA* architecture. First, *N* tiles are sampled from the WSI, and a feature vector is extracted from each tile using a ResNet-50 module pretrained on ImageNet. We then cluster the feature vectors into *K* clusters, and an average feature vector is obtained from each cluster, resulting in *K* aggregated feature vectors. Next, a transformer encoder and dense layers translate the obtained *K* feature vectors to predicted gene expression values. d) Performance of *SEQUOIA* compared to *HE*2*RNA*. For both architectures, we show the performance when trained from scratch and when finetuning from a model pretrained on normal tissues. Violin plots illustrate the distribution of Pearson correlation coefficients (left *y* axis) between the predicted and ground truth gene expression values in TCGA test sets. The top 1,000 genes with the highest correlation coefficients in each architecture are shown. Black squares indicate the absolute number (right *y* axis) of genes with significantly well-predicted expression levels. e) Distribution of RMSE values between the ground truth and predicted gene expression levels in TCGA test sets. WSI: whole-slide images; RMSE: root mean square error.

Since histological phenotypes and gene expression profiles vary across cancer types, the model was independently developed and validated in each cancer type. To evaluate the model, we carried out five-fold cross-validation. In each iteration, slides from 80% of the patients were allocated for training and internal validation, while the remaining 20% were reserved for testing. Prediction performance was assessed for each gene by comparing the predicted messenger RNA expression levels to the ground truth using Pearson’s correlation analysis and root mean squared error (RMSE). The resulting correlation coefficient and RMSE values were further compared to those obtained with a random, untrained model of the same architecture (see “Methods” for details). To identify genes with significantly well-predicted expression levels, we combined three criteria: (1) the predicted gene expression values must be significantly correlated with the ground truth, with a positive correlation coefficient and the associated *P* value smaller than 0.05 (*r*_1_ *>* 0 and *p*_1_ *<* 0.05) ; (2) *r*_1_ must be statistically higher than *r*_2_ (*r*_1_ *> r*_2_), as determined by Steiger’s Z test, where *r*_2_ represents the correlation coefficient obtained from the random model. For this comparison, we required the raw Steiger *P* value to be smaller than 0.05 (*p*_2_ *<* 0.05) and the adjusted *P* value by Benjamini-Hochberg correction smaller than 0.2 (*p*_3_ *<* 0.2); (3) The RMSE values obtained from the trained model must be smaller than those from the random model.

When trained from scratch, *SEQUOIA* was able to accurately predicted the expression levels of many genes. On average, 6,970 out of 25,749 genes were significantly well predicted across the nine cancer types (Figure 1d and Supplementary Table A4). Overall, the number of well-predicted genes was positively correlated with the number of training samples available in each cancer (Supplementary Table A4). The highest number (*N* = 12, 239) of genes was identified in BRCA, the cancer type with the most available slides (*N* = 1, 130 slides). Further, we identified 9,747 well-predicted genes in LUAD (*N* = 536 slides) and 6,985 genes in KIRC (*N* = 514 slides). Comparatively, PAAD and GBM had the lowest number of well-predicted genes as well as the lowest number of slides (PAAD: *N* = 418 genes from *N* = 202 slides; GBM: *N* = 4, 208 genes from *N* = 237 slides).

The correlation observed between the number of accurately predicted genes and the sample size suggests that the model can reach higher performance with a larger training dataset. Since published studies have revealed the advantages of pretraining transformer models on data of the same modality [22], we then pretrained the weights of transformer encoders using WSIs and RNA-seq data from normal tissues in the GTex cohort (A3) [27]. We found that finetuning a model pretrained on normal tissues increased the number of well-predicted genes in four cancer types (i.e., PAAD, KIRC, PRAD , COAD) (Figure 1d and Supplementary Table A4). Notably, incorporating the pretraining increased the number of significant genes by 8.1 times in PAAD, the cancer type with the least number of slides (*N* = 202 slides), and the median correlation coefficient increased by 1.5 times.

Since the histological appearance of BRCA has been shown to be associated with hormone receptor status [28], we separately assessed the performance in the estrogen receptor (ER) negative and ER positive subtypes. *SEQUOIA* accurately predicted the expression values for 8,517 and 3,840 genes in the ER positive and ER negative subtype, respectively (Supplementary Figure A1a). Of these genes, 2,103 genes were significantly predicted in both subtypes. These results demonstrate the capacity of *SEQUOIA* in predicting gene expression signals specific to breast cancer subtypes.

To compare the performance of our model with existing architectures, we benchmarked our results with the *HE*2*RNA* model [29]. *SEQUOIA* outperformed *HE*2*RNA* in all cancer types, irrespective of using the pretrained models or training from scratch (Figure 1d and Supplementary Table A4). On average, the number of genes with accurately predicted expression values increased by 8.2 times (7,464 genes versus 910 genes) with the pretrained *SEQUOIA* model compared to the counterpart of the *HE*2*RNA* model. Notably, in KIRP and LUAD, the number of accurately predicted genes increased by 23.5 and 14.6 times, respectively, with the *SEQUOIA* model. The cancer type with the smallest factor of increase (*×*1.9) was GBM, where *SEQUOIA* significantly predicted the expression levels for 2,498 genes compared to 1,303 genes using *HE*2*RNA*.

Finally, to compare the quality of predictions, we assessed both the correlation coefficients and RMSE values between the ground truth and predicted gene expression values obtained from the *SEQUOIA* and *HE*2*RNA* models. We found that *SEQUOIA* significantly outperformed *HE*2*RNA* in all cancer types (Figure 1d-e). On average, the median correlation coefficient was 2.0 times higher for *SEQUOIA* models compared to *HE*2*RNA*, and the RMSE value was 30% lower for *SEQUOIA* models (Figure 1d-e; Supplementary Tables A5 and A6; Supplementary Figure A2 and A3, Whitney U test, *P <* 0.0001).

### Pathway-level analysis of the predicted gene expression values

In our subsequent analysis, we focused on results obtained from *SEQUOIA* models pretrained on normal tissues. Genes with accurately predicted expression levels include protein-coding genes, long non-coding RNAs (lncRNAs) and micro-RNAs (miRNAs). On average, over 90% of the genes are protein-coding genes (Supplementary Figure A1b). To characterize their biological functions, we carried out gene set analysis. First, we performed hyper-geometric tests using the gene list with accurately predicted expression values in each cancer type. Second, we conducted gene set variation analysis (GSVA) using the predicted expression values and the pathway membership of each gene to assess the prediction accuracy at the pathway level [30].

In our hyper-geometric tests, we considered three gene set categories: (1) gene ontology, (2) KEGG pathway and (3) cell-type signatures. Gene ontology analysis revealed several common pathways enriched in the well-predicted genes across cancer types, including T cell activation (e.g., *CCL2*, *CCR2*, *CCDC88B* ), cell-matrix adhesion (e.g., *EMP2*, *COL16A1*, *VEGFA*), epithelial-mesenchymal transition (*BMP2*, *PDPN*, *SMAD2* ) and response to oxidative stress (e.g., *TP53*, *PRDX1*, *VRK2* ) (Figure 2a and Supplementary Data 1). Additionally, some gene sets were found enriched in specific cancer types. For instance, in GBM (Figure 2b) we identified genes associated with macrophage migration (*CCL2*, *EDN2*, *CKLF* ), protein kinase B signaling (*TNF*, *ADTRP*, *SETX* ), and endothelial cell development (*ROCK1*, *IKBKB*, *TNFRSF1A*). In LUSC (Supplementary Figure A1c), we identified genes associated with collagen biosynthetic process (*CREB3L1*, *WNT4*, *TGFB1* ), natural killer cell differentiation (*PTPRC*, *PIK3CD*, *KAT7* ), and histone H2A acetylation (*ACTL6A*, *MEAF6*, *DMAP1* ). Gene sets enriched in other cancer types can be found in Supplementary Data 1.

**Fig. 2:**
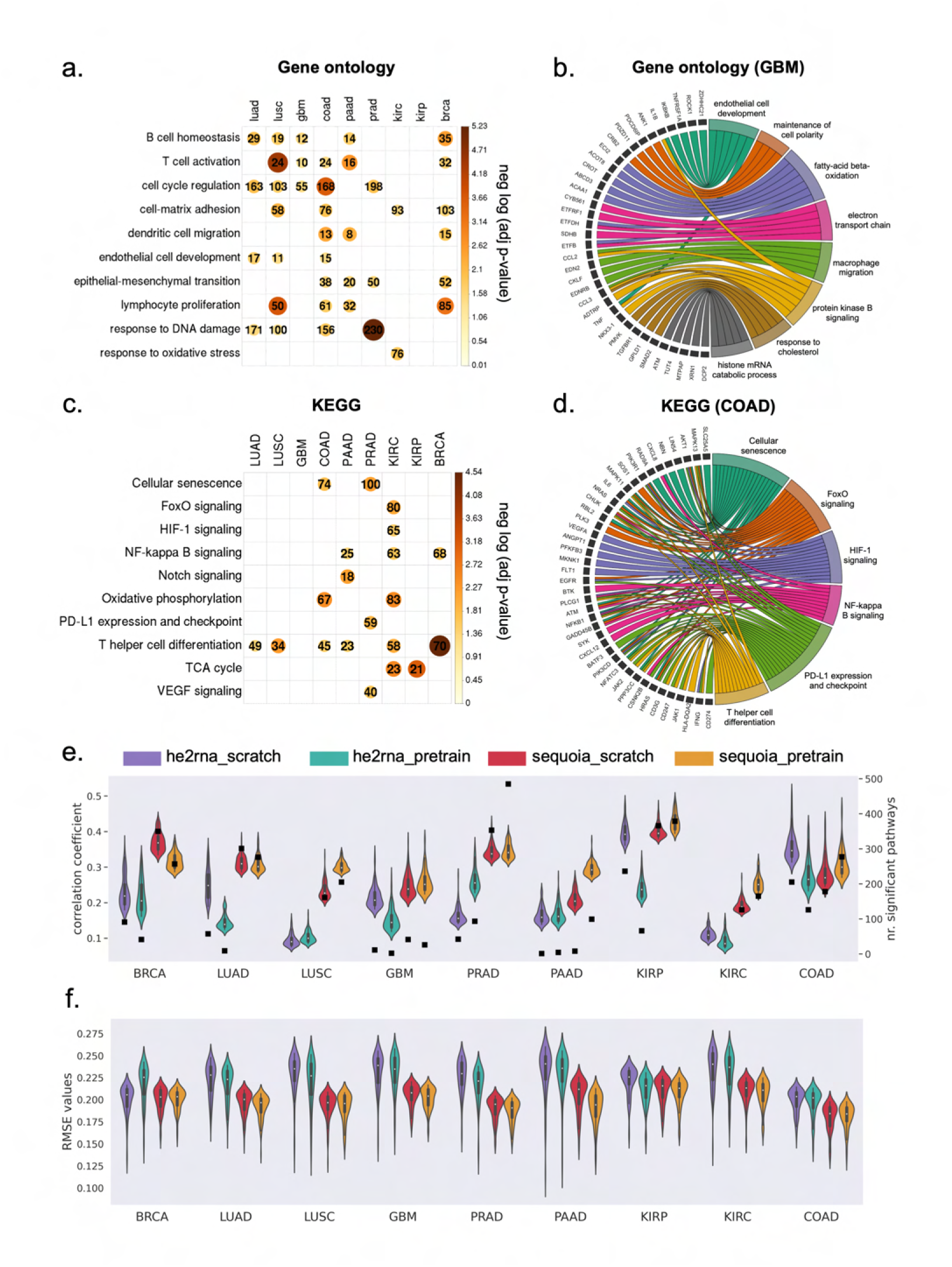
Evaluation of gene expression predictions at the pathway level. a) Heatmap showing significant *P* values obtained from hyper-geometric tests in gene ontology analysis of the well-predicted genes. Color and size of the circles represent the negative log-transformed *P* values. Integers represent the absolute gene count in each category, and non-significant categories are left in blank. b) Circos plot showing the enriched biological processes associated with the well-predicted genes in GBM. Gene names are displayed on the left and the corresponding biological processes are shown on the right. c) Heatmap showing the significant *P* values of the KEGG pathways across cancer types. Color and size of the circles represent the negative log-transformed *P* values. Integers represent the absolute gene count in each category, and non-significant categories are left in blank. d) Circos plot showing the KEGG pathways associated with the well-predicted genes in COAD. Gene names are displayed on the left and the corresponding pathways on the right. e) Violin plots illustrating the distribution of Pearson correlation coefficients (left *y* axis) between the predicted and ground truth pathway enrichment scores in TCGA test sets. The top 100 pathways with the highest correlation coefficients in each model are shown. Black squares indicate the absolute number (right *y* axis) of pathways with significantly well-predicted enrichment scores in each cancer. f) Violin plots illustrating the distribution of RMSE values between the predicted and ground truth pathway enrichment scores in TCGA test sets. The top 100 pathways with the lowest RMSE values in each model are shown. RMSE: root mean square error.

Furthermore, KEGG pathway analysis revealed that genes with accurately predicted expression levels are involved in the PD-L1 expression and check point pathway (*CD247*, *CD274*, *MAPK11* ), NF-kappa B signaling (*CXCL12*, *NFKB1*, *PRKCB* ), HIF-1 signaling (*GAPDH*, *HIF1A*, *VEGFA*), and VEGF signaling (*SPHK1, HRAS*, *HSPB1* ) (Figures 2c-d and Supplementary Data 2). Moreover, the well-predicted genes include cell-type markers for endothelial cells (*CD69*, *CD93* ), CD4 T cell (*CD3E*, *CD4*, *CD48* ), M2 macrophage (*CD14*, *CD163*, *CD84* ), and B cell (*CD19*, *CD53*, *CD37* ) (Supplementary Figure A1d and Supplementary Data 3). Overall, these results highlight the critical biological functions of the accurately predicted genes in regulating cell cycles, inflammation and hypoxia response.

To assess the prediction performance at the pathway level, we performed GSVA analysis[30]. An enrichment score was assigned to each gene ontology or KEGG pathway (total *N* = 3,120) based on the predicted expression values of individual genes and their pathway membership. The predicted enrichment score of each pathway was then compared to the ground truth using Pearson correlation analysis and RMSE values. To select significant pathways with accurately predicted enrichment scores, we compared the results to those obtained from the random model of the same architecture, using the same statistical methods and thresholds as we defined for selecting individual genes (Methods). On average, the enrichment score for 241 out of 3,120 pathways were significantly well predicted across the nine cancer types (Figure 2e). The accurately predicted pathways were similar to those identified using hyper-geometric tests, including pathways regulating cell cycle, inflammatory response and extracellular matrix organization.

When comparing the prediction performance between the *SEQUOIA* models pretrained on normal tissues versus models trained from scratch, the pretrained models achieved significantly higher correlation coefficients in six out of nine cancer types, including PAAD, LUSC, KIRC KIRP, COAD and GBM (Figure 2e, Mann-Whitney U test, *P <* 0.002). Of note, the number of accurately predicted pathways in PAAD was 12.3 times higher in the pretrained model compared to the scratch, and the median correlation coefficient was 1.4 times higher in the pretrained model (Mann-Whitney U test, *P <* 1*e −* 34). Moreover, in KIRC and LUSC, we observed an average of 1.4-time increase in median correlation coefficients using the pretrained model (Mann-Whitney U test, *P <* 4*e −* 33). These results indicated that pretraining the *SEQUOIA* models on data from normal tissues increased the prediction performance at pathway levels in specific cancer types.

When comparing the GSVA results obtained from the pretrained *SEQUOIA* models versus those from the *HE2RNA* models, we found *SEQUOIA* models achieved significantly higher correlation coefficients and lower RMSE values between the ground truth and the predicted pathway scores in all cancer types (Figure 2e-f, Mann-Whitney U test, *P <* 3*e −* 10). Overall, these results highlighted the advanced prediction performance of *SEQUOIA* models in predicting gene expression values at pathway levels.

### *SEQUOIA* generalizes to independent cohorts

Deep learning models trained on a specific dataset may be subject to bias due to technical noise (e.g., stain variations and color range), potentially leading to overfitting and limiting their ability to generalize to other datasets. To test the generalization capacity of *SEQUOIA*, we applied the models developed in the TCGA cohort to the matched cancer type in the CPTAC (Clinical Proteomic Tumor Analysis Consortium) cohort [3, 31–36]. We extended our validation to seven cancers from six tissues, including breast, lung, kidney, brain, colon and pancreas (Supplementary Table A2). For each gene, we compared the predicted expression levels in each cancer type to the ground truth using correlation coefficients and RMSE values as described for our TCGA analysis. This list of accurately predicted genes identified in the CPTAC cohort was then compared to the the gene list discovered in the TCGA cohort. Since the cohort size in CPTAC is smaller compared to the TCGA (Supplementary Table A2), it is expected that fewer genes can pass our significance thresholds. Despite this limitation, we were able to validate the prediction performance on many genes (Figure 3a). In BRCA, we validated the accuracy of predictions on 8,587 genes in the CPTAC cohort, which overlapped with 78% of the well-predicted genes (*N* = 11,069 genes) identified in the TCGA cohort. Additionally, we validated the accuracy of gene expression predictions for 7,259 (72%) genes in KIRC, 3,560 (72%) genes in LUSC, 5,330 (61%) genes in LUAD, 1,464 (59%) genes in GBM, 3,941 (51 %) genes in COAD and 1,177 (35%) genes in PAAD. Gene ontology analysis revealed the key functions of the validated genes in regulating angiogenesis, inflammatory response and cell cycle (Figure 3b). Furthermore, KEGG pathway analysis showed that the validated genes are associated with cell adhesion, NF-kappa B signaling, PI3K-Akt signaling and p53 signaling pathway (Figure 3c).

**Fig. 3:**
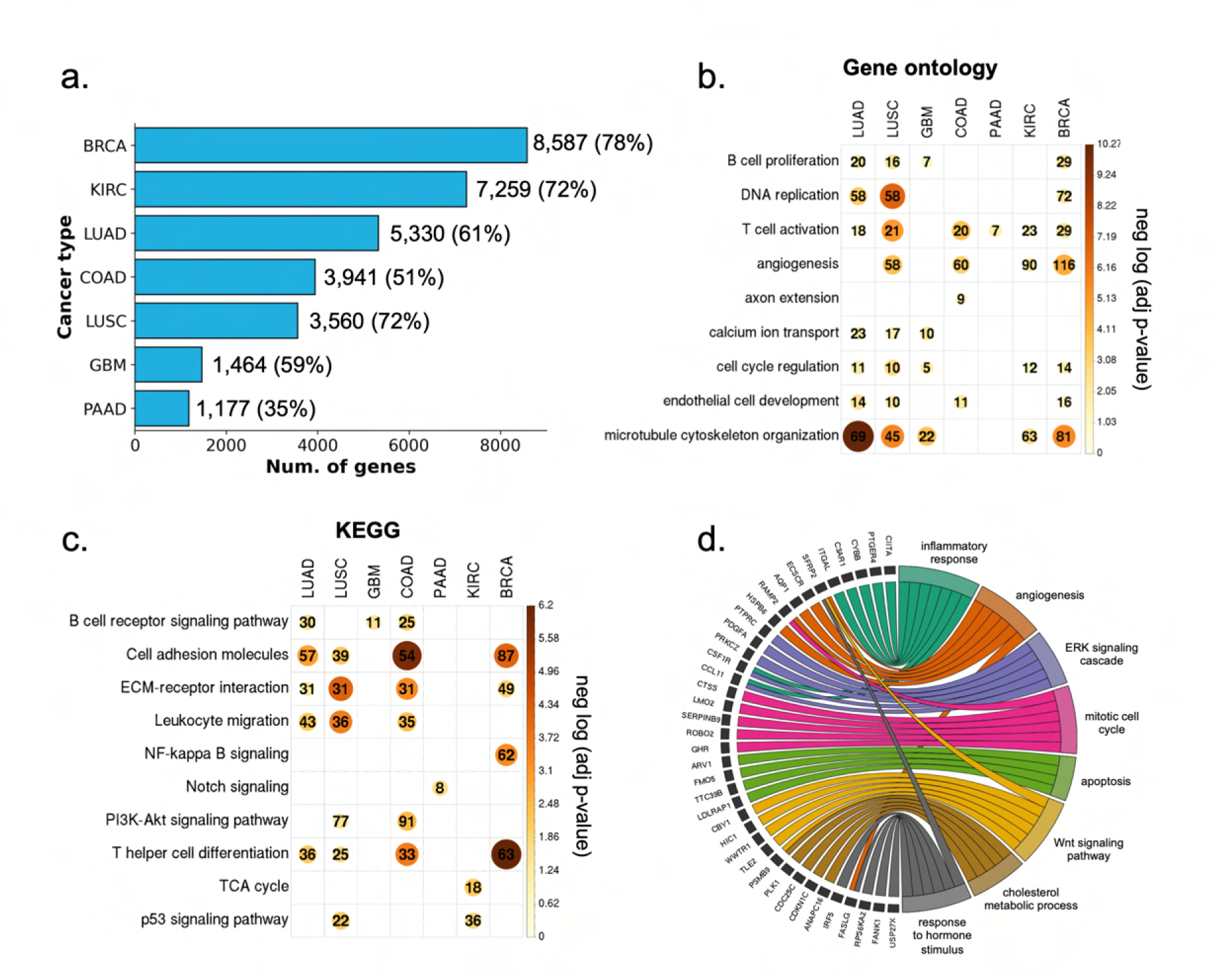
Characterization of the genes validated in external cancer cohorts. a) The number of genes validated in the CPTAC cohort. Percentages enclosed within the parenthesis indicate the proportion of significant genes discovered in the TCGA cohort that were validated in the CPTAC cohort. b) Heatmap showing the significant *P* values from the gene ontology analysis of the validated genes. Color and size of the circles represent the negative log-transformed *P* values. Integers represent the absolute gene count in each category, and non-significant categories are left in blank. c) Heatmap showing the significant *P* values from the KEGG analysis of the validated genes. Color and size of the circles represent the negative log-transformed *P* values. Integers represent the absolute gene count in each category, and non-significant categories are left in blank. d) Circos plot showing the enriched biological processes associated with the validated genes in lung adenocarcinoma.*P* values were adjusted for multiple testing using the Benjamini–Hochberg method.

To benchmark the generalization capacity of our model to existing architectures, we compared the prediction results to those from the *HE*2*RNA* model. Although *HE*2*RNA* identified numerous genes with accurately predicted expression values in both the TCGA and CPTAC cohorts, only a limited overlap of these genes was found between the two cohorts (Supplementary Table A7).

To further test the generalization capacity of our models, we extended the validation to a lung adenocaricnoma (LUAD) cohort from Tempus (“Methods”, *N* = 287 slides from *N* = 249 patients). This led to the identification of 1,217 genes that were well-predicted across all three (TCGA, CPTAC and Tempus) cohorts for patients with lung adenocarcinoma. Functional analysis of these genes revealed their regulatory functions in inflammatory response (*ITGAL*, *CYBB*, *PTGER4* ), angiogenesis (*VEGFD*, *TSPAN12*, *EMP2* ), ERK signaling (*PRKCZ*, *PDGFA*, *FGF10* ), and Wnt signaling pathway (*WIF1*, *SFRP2*, *NKD2* ) (Figure 3d). These results demonstrate the generalization capacity of *SEQUOIA* in predicting gene expression values across independent cohorts.

### A digital signature for breast cancer recurrence prediction

Given that *SEQUOIA* was able to predict the transcriptional activity of genes involved in key cancer-related pathways (Figures 2 and 3), we next assessed whether these genes have prognostic value. We focused our analysis on breast cancer, in which the highest number of genes (*N* = 11,069) were accurately predicted for their expression levels.

The accurately predicted genes encompass various published prognostic signatures (Supplementary Data 5) [37–40]. These include 47 out of 50 (94%) genes of the PAM50 signature, all 12 (100%) genes of the EndoPredict signature, 13 out of 21 (62%) genes of the Oncotype DX signature, 47 out of 70 (67%) genes of the Mammaprint signature, 6 out of 7 (86%) genes of the Breast Cancer Index and 4 out of 5 (80%) gene of the Mammostrat signature.

Since the Oncotype DX and MammaPrint signatures are developed on data from RT-qPCR and microarray assays, and their exact mathematical formulas are protected by proprietary license, we next sought to develop a RNA-seq-based gene expression signature that can stratify the risk of breast cancer recurrence. We fitted a regularized Cox regression model on the ground-truth gene expression values for the accurately predicted genes from *SEQUOIA*, and the regression model aims to predict a risk score of recurrence for each patient (see ”Methods” for details). High risk scores indicate a greater likelihood of recurrence. The model was developed on the TCGA cohort (*N* = 858 patients) and independently validated using data from the ”SCANB” cohort (*N* = 5,034 patients). Moreover, to test whether our signature can be generalized to traditional microarray-based gene expression data, we further validated it using an independent dataset of the ”METABRIC” cohort (*N* = 2,262 patients) [41].

Our analysis led to the identification of a 50-gene signature significantly associated with recurrence (Figures 4a-c and Supplementary Data 6). To assess its performance, we first treated the predicted risk score as a continuous variable. Results from univariate Cox regression analyses (Figures 4a-b and Supplementary Figure A1e) showed that the predicted risk scores were significantly associated with recurrence-free survival: TCGA (HR = 2.24, .95CI = 1.99-2.52, *P <* 2*e −* 16), SCANB (HR = 1.31, .95CI =1.25-1.39, *P <* 2*e −* 16), and METABRIC (HR = 1.21, .95CI = 1.08-1.36, *P* = 7.8*e −* 07). To further assess the model, we treated the predicted risk score as a dichotomous variable. Patients within each cohort were divided into a high-risk and a low-risk group based on the median risk score (Figures 4a-b and Supplementary Figure A1e). Results from the log-rank test demonstrate that the high-risk group had significantly worse prognosis compared to the low risk group: TCGA (*P <* 2*e −* 16), SCANB (*P* = 4*e −* 13), and METABRIC (*P* = 8*e −* 05).

**Fig. 4:**
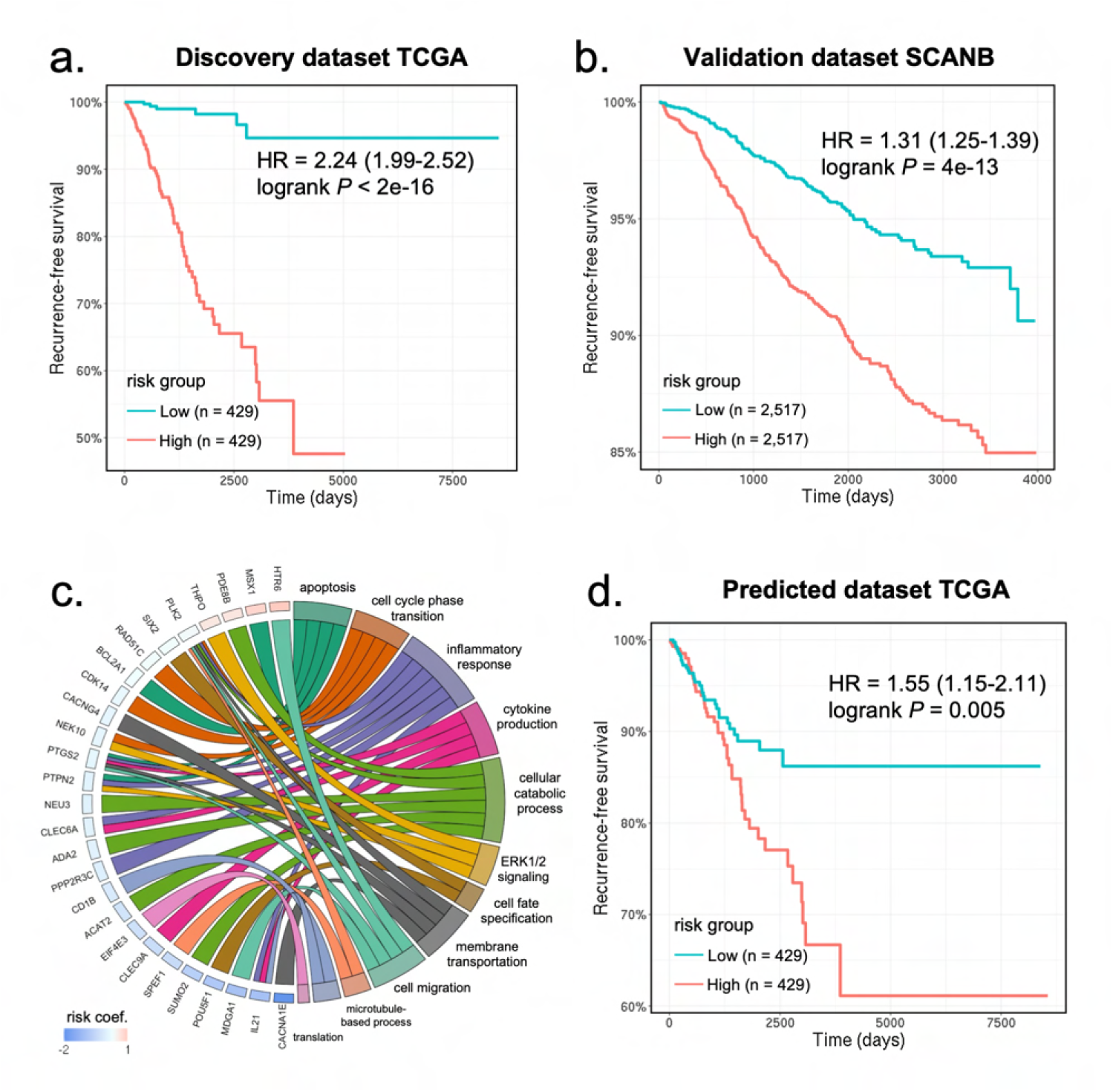
Development and validation of a digital signature for predicting breast cancer recurrence. a) Kaplan-Meier curves of recurrence-free survival obtained from the TCGA discovery dataset. Patients were split by the median risk score. b) Kaplan-Meier curves of recurrence-free survival in the SCANB validation dataset. c) Circos plot showing the biological processes associated with the prognostic gene signature. Gene names and the associated risk coefficients are shown on the left and the corresponding biological processes are shown on the right. d) Kaplan-Meier curves of recurrence-free survival obtained from the predicted gene expression values in the TCGA dataset. Patients were split by the median risk score. HR: hazard ratio.

To assess whether breast cancer subtype was a confounding variable in risk prediction, we incorporated the PAM50 molecular subtypes and hormone (i.e., estrogen and progesterone) receptor status as covariates into our Cox regression analyses. We found that the predicted risk score was still significantly associated with prognosis after including these covariates: TCGA (HR = 2.22, .95CI =1.96-2.52, *P <* 2*e −* 16), SCANB (HR = 1.26, .95CI =1.20-1.33, *P <* 2*e −* 16), METABRIC (HR = 1.22, .95CI = 1.08-1.39, *P* = 7.6*e −* 05).

Gene ontology analysis (Figure 4c) revealed the regulatory functions of the signature genes in cell apoptosis (*MSX1*, *PTPN2*, *BCL2A1* ), cell-cycle phase transition (*NEK10*, *CDK14*, *RAD51C*, *PLK2* ), inflammatory response (*IL21*, *PTGS2*, *PPP2R3C* ), cytokine production (*CLEC9A*, *CLEC6A*), cellular metabolic process (*PDE8B*, *ADA2*, *NEU3*, *SUMO2*, *ACAT2* ), ERK signaling cascade (*THPO*, *PTPN2* ), cell-fate specification (*POU5F1*, *SIX2* ), cell membrane transportation (*CACNA1E*, *PTGS2*, *CACNG4*, *PLK2* ), and cell migration (*MDGA1*, *HTR6* ).

So far, we have developed and validated a 50-gene signature using the ground-truth gene expression values. We then tested whether utilizing the gene expression values predicted from histology images alone was sufficient to stratify the risk groups. For each patient, we calculated a risk score using the same risk coefficient in our Cox regression model, but this time replacing the ground-truth gene expression values with the predicted values. As shown in Figure 4d, patients assigned with high risk scores demonstrated significantly worse prognosis compared to patients with low risk scores (Cox regression: HR = 1.55, .95Cl = 1.15-2.11, *P* = 0.006; Log-rank test: *P* = 0.005). These results indicate that *SEQUOIA* can accurately predict the expression of genes associated with breast cancer recurrence.

### Tile-level predictions validated with spatial transcriptomics

So far, we have demonstrated the ability of *SEQUOIA* in predicting RNA-Seq gene expression values collected from bulk tissues. However, gene expression patterns are known to vary across different tumor regions due to intra-tumoral heterogeneity that results from uneven spatial distributions of cell phenotypes. Uncovering spatial gene expression patterns can reveal the intricate landscape of tumor architecture and signaling environment, which is known to affect tumor growth, metabolic processes, and resistance to therapy [6, 8]. We hence investigated whether our models trained at the slide-level can be used to predict gene expression values at locoregional levels within tumor tissues.

Here, we implemented a sliding-window approach to generate tile-level predictions of gene expression. The histology image was first processed using a sliding window of 10*×*10 tiles starting from the left upper corner, where the dimensions of each tile (128*um×*128*um*) were consistent with those used during the training of the *SEQUOIA* models. For each window, a 100 *×* 2048 feature vector was extracted and used as input to *SEQUOIA* for generating a prediction. This prediction was then stored for every tile within the window (see “Methods” for details). After processing the entire image, the predicted gene expression for each tile was calculated as the average of the stored values for that tile. A stride of 1 (tile) was chosen for fine-grained analysis. To validate the prediction, we utilized data from an independent cohort of patients with glioblastoma (GBM), which contains matched histology images and spatial transcriptomics data (*N* = 54, 000 gene expression spots from *N* = 18 patients), providing tile-level ground truth gene expression measurements [5] (Figure 5a). We focused our analysis on the top 500 genes for which *SEQUOIA* generated the best predictions on the TCGA test set (i.e. genes with the highest Pearson correlation coefficients). For each of these genes, we generated a spatial heatmap illustrating their expression values across the slide. To quantitatively assess the prediction performance, we used the Earth Mover’s Distance (EMD) as an evaluation metric (“Methods”). EMD values are bounded between 0 and 1, with lower values indicating a closer correspondence between predictions and ground truth. On average, *SEQUOIA* achieved an EMD of 0.15 (.95CI = 0.148-0.152) across all slides and genes (Supplementary Table A8). Higher performance was observed in slides with high degrees of spatial variance in gene expression [5, 6].

**Fig. 5:**
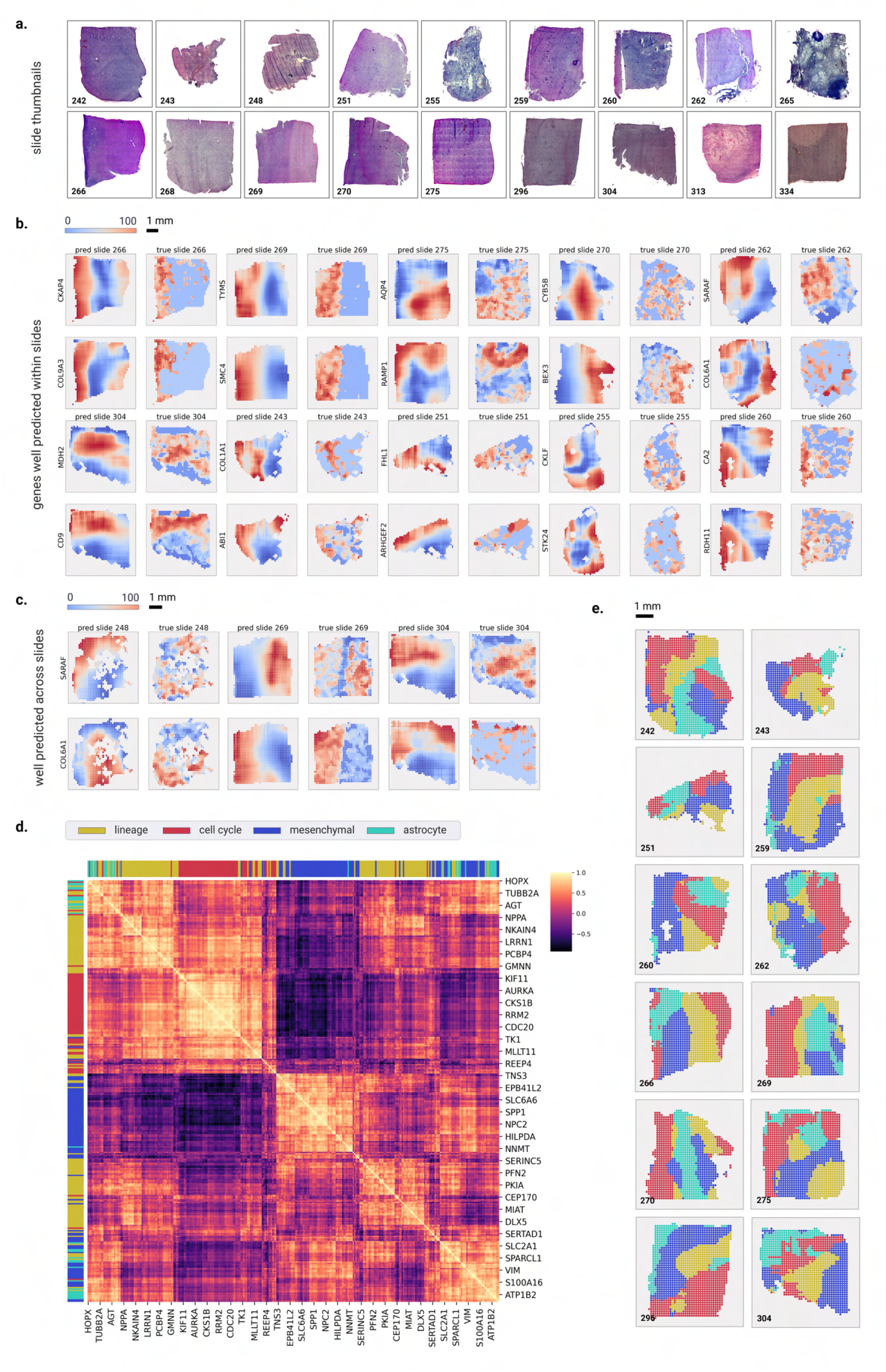
Spatial visualization of gene expression predicted at the tile level. a) Whole Slide Image thumbnails from the validation cohort. b) Examples of genes that are well-predicted spatially within slides, with predicted spatial gene expression on the left and ground truth on the right. The prediction and ground truth maps were normalized to percentile scores between 0-100. c) Examples of genes that are spatially well-predicted across several slides. Each row shows the prediction map (on the left) and ground truth (on the right) for a particular gene across four slides. d) Heatmap showing the correlation coefficients of meta-gene modules that define the transcriptional subtype and proliferation state of GBM cells. e) Spatial organization of the predicted transcriptional subtypes within different slides. Transcriptional subtypes were assigned based on the meta-gene module showing the highest prediction values

Notably, *SEQUOIA* generated accurate spatial predictions for genes that hold significant relevance to GBM malignancy and prognosis. For instance, the *COL6A1* and *COL9A3* genes are highly expressed in the mesenchymal subtype of GBM, a subtype associated with unfavorable prognoses [42, 43]. In our spatial predictions, we observed a median EMD of 0.11 across all slides for both *COL6A1* and *COL9A3* (Figures 5b-c). Furthermore, the *CKAP4* gene, with and EMD of 0.12 has been shown to mediate the growth, migration, and invasion of GBM cells [44]. These results highlight the potential of *SEQUOIA* in accurately predicting spatial gene expression patterns related to GBM malignancy and prognosis.

The integrative analysis of single-cell RNA-seq and spatial transcriptomics data in recent studies have revealed that cells sharing the same transcriptional subtype are often co-localized within spatially segmented niches [5, 45]. To investigate whether *SEQUOIA* captured true biological signals that reflect underlying tissue compositions, we assessed spatial co-expression patterns of functionally related genes. We considered four previously established meta-gene modules governing the transcriptional subtype and proliferation state of GBM cells: (1) ‘lineage development’ (124 genes), (2) ‘cell cycle’ (70 genes), (3) ‘mesenchymal-like’ (92 genes) and (4) ‘astrocyte-like’ (37 genes) [46]. Spatial correlation analyses showed that genes within the same meta module consistently clustered together, exhibiting similar spatial expression patterns (Figures 5d-e). To demonstrate the spatial prediction capacity of *SEQUOIA* in other cancer types, we developed an user-friendly, interactive web application (https://sequoia.stanford.edu) where users can explore the spatial heatmap for genes predicted in the TCGA cohorts. These results demonstrate the potential of *SEQUOIA* in resolving spatial cellular architectures within heterogeneous tumor tissues.

## Discussion

Transcriptomic analysis of tumor tissues holds immense promise in advancing personalized diagnosis and outcome predictions. In this study, we presented *SEQUOIA*, a deep learning model for predicting RNA-seq gene expression data from Whole Slide Images (WSIs). We combined algorithmic and methodological advancements, followed by thorough analyses of gene functions, clinical relevance, and generalization capacity. Through a comprehensive evaluation of our model in nine cancer types across seven tissues, we demonstrated the value of *SEQUOIA* in predicting clinically relevant gene expression patterns.

Over the past decade, deep learning has revolutionized cancer diagnosis. Published studies have demonstrated the potential of deep neural networks in extracting intricate patterns from medical images. He et al. developed ST-Net, a convolutional neural network that predicts the expression values of 250 genes from histology images in breast cancer [47]. Their model however is trained on individual tiles, which requires high-resolution training labels obtained from spatial transcriptomics assays and did not integrate contextual information across tiles. To model contextual information, Graziani et al. incorporate an attention mechanism into their model for gene expression predictions. However, this strategy requires training a dedicated model for predicting the expression of each individual gene [48]. While this approach reaches publishable performance, it can lead to computational challenges, particularly when attempting to infer the entire transcriptome. In a recent study, Alsaafin et al. utilized transformer modules to extract latent representation of WSIs. Their model, trained exclusively in renal cell carcinoma, was used for gene expression predictions and subtype classifications [25]. However, the generalization capacity of the model to other cancer types remain to be determined. Moreover, published studies have indicated that transformer-based models outperforms convolutional neural network only when a pretraining step using large dataset is incorporated[26]. Moreover, the increased complexity of transformer-based models may lead to overfitting, especially with limited training data[49].

To address these challenges, we pretrained the weight parameters of the transformer encoder using data from normal tissues. Our results showed that the pretraining regimen improved the prediction performance at both individual gene levels and at the pathway level. The observed improvement was most significant in cancers with small training datasets. To demonstrate the advantages of our model, we compared it with *HE*2*RNA* [29], a recent model for whole transcriptome prediction from WSIs. The results of our analysis revealed consistent improvements across various cancer types when using *SEQUOIA* in comparison to *HE*2*RNA*. A key factor driving this performance boost lies in the attention-based mechanism leveraged by *SEQUOIA*, which enables effective integration of information between tiles. In contrast, *HE*2*RNA* treated each tile as an independent entity, limiting its ability to capture the contextual relationships present in the data.

The genes with accurately predicted expression values by *SEQUOIA* were found to be associated with key pathways pertinent to cancer progression. Among these were genes involved in regulating cell cycles, inflammation, angiogenesis, and hypoxia response. Additionally, the model effectively captured cell-type markers, including those for endothelial cells, CD4 T cells, M2 macrophages and B cells. Building upon the well-predicted genes, we developed a 50-gene signature that predicts the risk of breast cancer recurrence. Although the gene expression signature was developed on ground-truth gene expression values, we demonstrated its utility in patient stratification by just using the predicted gene expression. Despite the decreasing costs for transcriptomics sequencing, the integration of gene expression analysis into clinical routines is hindered by the lack of necessary equipment and trained personnel. By leveraging *SEQUOIA*’s predictions, one can gain mechanistic insights linking histopathological phenotypes to molecular characteristics, thereby offering guidance for disease classification, prognostication, and treatment planning.

Understanding spatial topological organization of tumor cells has attracted recent research interest in the field. The advancement of spatial transcriptomics and proteomics technologies have deepened our understanding of the intrinsic signaling environment that drive tumor growth, metastasis and treatment sensitivity. While *SEQUOIA* was trained using bulk RNA gene expression, we demonstrated its potential in predicting gene expression patterns at the locoregional level. We implemented a technique that enables computational reconstruction of high-resolution spatial gene expression within tumor tissues. The results were validated using a spatial transcriptomics dataset obtained from an independent cohort of glioblastoma patients. Notably, genes with accurate spatial expression predictions include those regulating malignant phenotype and prognosis. Applying such computational method to WSIs can bring significant values to both clinical and research settings. In the clinic, it can aid in identifying specific regions within a heterogeneous tumor that require sequencing, hence ensuring the accurate detection of biomarkers and preventing the omission of critical lesions [18]. In research, this approach enables the cost-efficient exploration of gene expression dynamics at high resolution, which allows to generate hypotheses about signaling events driving cellular interactions, thereby advancing our understanding of the complex mechanisms underlying cancer progression.

In the future, the accurate prediction of molecular traits from histology holds immense potential for enhancing diagnosis and prognosis for cancer. The predicted gene expression and pathway activation levels can provide valuable insights into a tumor’s aggressiveness and its molecular characteristics, advancing our understanding of cancer heterogeneity and enabling personalized and targeted therapies. Once clinically validated and further improved (e.g. by training on large, multi-center cohorts), the implementation of such predictive models has the potential to streamline medical processes, save costs, and improve efficiency by rapidly identifying actionable information from image-based data.

In conclusion, by combining algorithmic advancements with thorough analyses of biological functions, clinical relevance, and generalization capacity, our research demonstrates the potential of using transformer-based deep learning models in predicting high-dimensional gene expression features from whole-slide histology images.

## Methods

### Patient cohorts and ethics

#### TCGA

For model training and evaluation, anonymized patient data were retrieved from the publicly available The Cancer Genome Atlas (TCGA) archive (available at https://portal.gdc.cancer.gov). We used paraffin-embedded (FFPE) whole slide images (WSIs) and matched gene expression data of nine cancer types, including prostate adenocarcinoma (PRAD), pancreatic adenocarcinoma (PAAD), lung adenocarcinoma (LUAD), lung squamous cell carcinoma (LUSC), kidney renal papillary cell carcinoma (KIRP), kidney renal clear cell carcinoma (KIRC), glioblastoma multiforme (GBM), colon adenocarcinoma (COAD), and breast adenocarcinoma (BRCA). The number of patients, WSIs and genes used for training each cancer type is listed in Supplementary Table A1.

#### CPTAC

For validation, anonymized patient data were retrieved from the publicly available Clinical Proteomic Tumor Analysis Consortium (CPTAC) cohort (https://portal.gdc.cancer.gov). We downloaded matched WSIs and gene expression data from seven cancer types from six tissues, including breast invasive carcinoma (BRCA), lung adenocarcinoma (LUAD), lung squamous cell carcinoma (LSCC/LUSC), colon adenocarcinoma (COAD), kidney renal clear cell carcinoma (CCRCC/KIRC), glioblastoma multiforme (GBM), pancreatic adenocarcinoma (PDA/PAAD). The sample size is described in Supplementary Table A2.

### Tempus

For an additional validation, we utilized matched WSIs and RNA-seq data (*N* = 287 slides from *N* = 249 patients) of lung adenocarcinoma (LUAD). The data were obtained through a data transfer agreement with Tempus Labs, Inc.

### GTex

For pre-training the transformer encoder, we used WSIs and gene expression data from six normal tissues (i.e., brain, colon, kidney, lung, pancreas, prostate). Data were obtained from The Genotype-Tissue Expression (GTEx) project (https://gtexportal.org), and the sample size is described in Supplementary Table A3

### Spatial GBM, SCANB, METABRIC

Spatial transcriptomic data and matched histology images of GBM were obtained from a published study by Ravi et al. (https://datadryad.org/stash/dataset/doi:10.5061/dryad.h70rxwdmj) [5]. Data of the SCANB and METABRIC breast cancer cohorts were obtained from published studies by Staaf et al.[50] and Curties et al.[41].

### Preprocessing of RNA-Seq data

For training and validation of our models, we used FPKM-UQ normalized gene expression values. Since the gene expression values span several orders of magnitude and our model was trained using the Mean Squared Error loss function, the training process may introduce bias to genes with large gene expression values. To overcome this potential bias, we performed log2 transformation (*v → log*_2_(*v* + 1)) of the gene expression values.

For pre-training the transformer encoder, we obtained the RNA-seq data of normal tissues from the GTEx data portal (https://gtexportal.org/home/datasets). We performed the same log2 transformation (*v → log*_2_(*v* + 1)) of the gene expression values. Since during the pre-training phase, we combined data from all tissue types, we performed a *z* -score normalization of the gene expression values in each individual tissue type, and the normalized gene expression matrices were concatenated across the tissue types.

We focused our analysis on three gene categories: (1) protein-coding genes, (2) micro-RNAs (miRNAs) and (3) long non-coding RNAs (lncRNAs). On average, the protein-coding genes account for 85% of all the analyzed genes.

### Preprocessing of Whole Slide Images

Whole-slide images (WSIs) were acquired in *SVS* format and downsampled to 20*×* magnification (0.5*µm* px^-1^). We used the Otsu threshold method to obtain a mask of the tissue, which allows to omit tiles mostly containing white background [51]. WSIs have much larger dimensions than natural images (usually over 10*k ×* 10*k* pixels), and therefore cannot be used directly to train machine learning models. To address this challenge, we employed a multiple instance learning (MIL) approach, where each WSI was cropped into non-overlapping tiles of 256 *×* 256 pixels (128*µm ×* 128*µm*). In each slide, we randomly selected a maximum of *N* = 4000 tiles, omitting those containing more than 20% background and tiles with low contrast.

To obtain a feature representation at slide-level, we organized the selected tiles into bags. First, we used a ResNet-50 module pre-trained on ImageNet to covert each tile into a feature vector (1 *×* 2048). Then, we used the k-means algorithm to cluster similar tiles of each slide into *K* = 100 clusters. Each cluster contains tiles with similar morphological features, where cluster *A* may represent tiles that mostly contain tumor cells, cluster *B* may contain tiles with mostly connective tissue and so on. Features of patches within the same cluster were averaged, resulting in a matrix of 100 *×* 2048 vectors that represent the slide.

### *SEQUOIA* architecture

*SEQUOIA* is inspired by the vanilla Vision Transformer (ViT) architecture [26] which extrapolates the Transformer architecture from the natural language processing (NLP) domain to computer vision [52]. For a ViT, an image is divided in patches of 16 *×* 16 px, representing “tokens” of the image. Feature vectors are then extracted from these patches by linear projection in the ViT. Then, they are fed to a transformer encoder, which outputs a new representation of the input and forwards it to a multi-layer perceptron (MLP) head that makes the final prediction.

In our work, the division of an image into small patches corresponds to the WSI being divided into the *K* feature clusters (*K* = 100). The 100 *×* 2048 feature matrix is fed to the transformer encoder, which comprises 6 encoder blocks, 16 attention heads, and a head dimension of 64. The transformer encoder allows modeling the relationship across feature clusters before deciding whether they are relevant for the slide-level prediction. After layer normalization, the output is sent to an MLP layer with dimension 2048 *× num genes*, with *num genes* the number of genes available to predict in each cancer type (see Supplementary Table A1).

### Pretraining on normal tissues

We performed pre-training of the *SEQUOIA* and *HE*2*RNA* models using normal tissues corresponding to the tested cancer types (Supplementary Table A3). We combined data from all normal tissues for the pre-training step, and the resulting model was afterwards finetuned for each specific cancer type.

Notably, in our preliminary experiments, we observed that incorporating breast normal tissues into the pre-training led to an overall decrease in performance compared to the model pre-trained on all other normal tissues without the breast. This decrease can likely be attributed to the known differences in tissue composition between the normal breast and breast cancer. While the normal breast mainly comprises adipocytes, the tumors consist of transformed epithelial cells. Consequently, we excluded normal breast tissue from the pre-training process, resulting in six tissue types used for pre-training: lung, brain, kidney, pancreas, prostate, colon. We utilized the gene expression data for 19,198 genes available across all normal tissues.

The model was trained using the Mean Squared Error loss function for 200 epochs with early stopping (i.e., early stop if the loss did not decrease for a *patience* of 100 epochs), and a batch size of 16. Model parameters were optimized using the Adam optimizer with a learning rate of 3 *×* 10*^−^*^3^.

### Training details

After pretraining the model on normal tissues, we finetuned a dedicated model for each cancer type using data of the TCGA cohort. The transformer encoder was initialized with the pre-trained weights, and a new prediction head, consisting of a layer norm and a linear layer, was trained to predict the gene expression levels in cancer tissues (see Figure 1).

For training and evaluation of the model, we conducted a five-fold cross-validation using data from the TCGA cohort. In each fold *i*, the dataset was partitioned on patient level, allocating 80% for ‘global‘ training and 20% for testing. To determine the optimal stop point for training the model in fold *i*, the ‘global‘ training set *i* was further split into 90% for training and 10% for internal validation. We used the Mean Squared Error (MSE) as the loss function during model training, with each model being trained for a maximum of 200 epochs. For early stopping and determining the point for model saving, instead of relying solely on the Pearson correlation coefficient as described in Schmauch et al. [29] , we employed a criterion that considers both MSE and correlation. Specifically, we continued training and saved the model at each optimal MSE point as long as the MSE continued to decrease. Once the MSE stopped improving for a consecutive *patience* interval of 100 epochs, we continued the training process if the correlation had improved in the last *patience* epochs and if the MSE remained below a reasonable threshold (i.e. *MSE < δ* + *bestMSE, with δ* = 0.5). We then saved the model at the optimal epoch if the correlation had improved (i.e. *corr > best corr*). Throughout this process, we used a fixed learning rate of 1*×*10*^−^*^3^ and batch size of 16, and the model parameters were optimized with the Adam optimizer.

### Identification of significantly well-predicted genes and pathways

To assess the performance within the TCGA cohort, we concatenated the predictions of all test sets *i* (*i* = 1..5). For each gene, the predicted gene expression values were compared to the ground truth using both Pearson’s correlation analysis and root mean squared error (RMSE). The resulting correlation coefficient and RMSE values were then compared to those obtained with a random, untrained model of the same architecture. To identify genes with significantly well-predicted expression levels, we combined three criteria: (1) The correlation coefficient (*r*_1_) between ground truth and the predicted gene expression values must be positive and the associated *P* value (*p*_1_) should be less than 0.05 (*r*_1_ *>* 0 and *p*_1_ *<* 0.05); (2) *r*_1_ must be significantly higher than *r*_2_ (*r*_1_ *> r*_2_) as determined by the Steiger’s Z test, where *r*_2_ represents the correlation coefficient between ground truth and predicted gene expression values obtained from the random model. We required the raw Steiger *P* value to be less than 0.05 (*p*_2_ ¡ 0.05) and the adjusted *P* value by Benjamini-Hochberg correction to be less than 0.2 (*p*_3_ ¡ 0.2); (3) The RMSE values obtained from the trained model must be smaller than those from the random model.

### Gene set analysis

The gene set analysis was performed with the ClusterProfiler R library (version 4.2.1) [53] and GSEApy package (version 1.0.5) [54]. Biological processes from gene ontology and cell-type signatures were obtained from the MSigDB database (https://www.gsea-msigdb.org/gsea). KEGG (Kyoto Encyclopedia of Genes and Genomes) pathway annotations were obtained from the KEGG database (https://www.genome.jp/kegg/catalog/orglist.html). The enrichment analysis was performed with hyper-geometric testing, and the *P* values were corrected with the Benjamini-Hochberg procedure. To generate heatmaps of the *P* values, we aggregated gene sets with high similarities (e.g., “regulation of T cell proliferation” and “positive regulation of T cell proliferation”), and the average *P* values were shown. Gene set variation analysis (GSVA) was performed to assign an enrichment score to each gene ontology or KEGG pathway based on the ground truth or predicted gene expression values using the GSEApy package (version 1.0.5).

### Identification and validation of the prognostic gene signature

To construct a gene expression model for predicting breast cancer recurrence, we first selected the top 5,000 well-predicted protein-coding genes from the TCGA-BRCA cohort as potential candidates. Then, we performed LASSO Cox regression model analysis with the ‘glmnet’ R package (version 4.1)[55]. The penalized Cox regression model with LASSO penalty was used to achieve shrinkage and variable selection simultaneously. The optimal value of the penalty parameter *λ* was determined through a five-fold cross-validation.

Utilizing the optimal *λ* value, we curated a list of prognostic genes, each associated with a coefficient (i.e., hazard ratio) that was not equal to zero. The risk score was derived by performing a linear combination of the expression levels of the selected genes, with each expression level being weighted by its associated coefficient, as described by the equation 1:

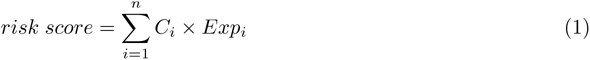

where *C_i_* represents the coefficient of a gene and *Exp_i_* its expression value.

The patients in each dataset were split into a low-risk and a high-risk group according to the median risk score. Finally, the Kaplan–Meier estimator and the log-rank test were performed to assess the difference in recurrence-free survival between the low-risk and high-risk groups.

### Spatial visualization of predicted gene expression on tile level

To visualize predicted gene expression spatially on tile-level, we implemented a sliding-window method. Starting from the left upper corner of the WSI, consider a window of 10 *×* 10 tiles. For clarity, we refer to the location of the window by the (*x, y*) coordinate of the left upper tile in the window. Hence, the window in the left upper corner has coordinate (0, 0) and is referred to as *w*_0,0_ (*x, y* axes defined as in image processing, origin in left upper corner and x-axis increases when moving to the right, y-axis increasing when moving below).

The 100 *×* 2048 feature vectors of tiles in the window *w_x,y_* are fed to the model (each individual tile feature vector serves as a ‘cluster mean vector’). The resulting predicted gene expression *g_w__x,y_* is saved for all tiles in the window. Then, the window is moved *stride* number of tiles to the right (*w_x_*_+_*_stride,y_*), and the predicted gene expression is again saved for each tile in the window. When the window has reached the end of a row (*x* + *stride* + 10 equals the width of the image), a new window is started at position *stride* below the previous row (*w*_0_*_,y_*_+_*_stride_*). After the window has passed the entire WSI, the prediction for each tile is calculated as the average of all values that were saved for that tile when it was part of a window *w_x,y_*. In our implementation, we chose *stride* = 1 (larger strides require less compute time but are less fine-grained).

For comparison of the predicted spatial gene expression with the spatial transcriptomics measurement in the ground truth, we resampled the ground truth resolution to match the predicted resolution. Namely, the ground truth resolution was 55*µm* per spot which is higher than the predicted resolution of 256*µm* per spot. Hence, we compared each spot in the prediction with the average of the four nearest spots in the ground truth (nearest in terms of smallest Euclidean distance between the x,y coordinates of the spots). We also performed median filtering on the ground truth map to remove noise (window size 3 *×* 3) and we only considered genes with *>*= 10 unique measured values in the spatial ground truth map (to avoid incorporating noisy measurements). Finally, we converted both the predicted and ground truth values to normalized percentile scores between 0-100.

### Earth Mover’s Distance

For a quantitative evaluation of the spatial visualization capabilities of the model, we used the two dimensional Earth Mover’s Distance (EMD) (implemented with the *cv*2*.EMD* function from opencv-python [56]). Intuitively, the metric captures the minimum amount of ‘work’ required to transform one distribution into the other. Often the two distributions are informally described as different ways of piling up earth/dirt, and the ‘work’ to transform one distribution into another is defined as the amount of dirt multiplied by the distance (Euclidean distance in our case) over which it is moved. Hence, this metric takes into account the *spatial context* to determine how well the prediction map corresponds to the ground truth. This is in contrast to pixel-level metrics which only take into account how correct a certain pixel is irrelevant of its 2D location and context (e.g. calculating for each pixel the Mean Squared Error between prediction and ground truth).

### Spatial correlation analysis of GBM signature genes

To assess whether genes exhibiting similar spatial expression patterns are functionally related, we used four recurrent meta-gene modules governing the transcriptional subtype and proliferation state of GBM cells as discovered from a published single-cell RNA-seq study [46]. We included all signature genes from these modules, except for those (*N* = 18 genes) not included in our training process. The neuralprogenitor-like (NPC-like) and oligodendrocyte-progenitor-like (OPC-like) modules were combined into one group, namely ’lineage development’, which includes a total of 124 genes. Further, gene modules regulating G1/S and G2/M phase transitions (*N* = 70 gens) were combined into a ’cell-cycle’ module. Finally, the ‘mesenchymal-like’ (*N* = 92 genes) and ‘astrocyte-like’ (*N* = 39 genes) modules were included as separate groups.

To assess spatial co-expression patterns, we determined the similarity of spatial prediction maps for each pairwise combination of genes (*N* = 325 genes in total). This was accomplished by first flattening the tile-level predictions into two 1D arrays and then computing the Pearson correlation between them. This process was repeated in each slide, and the resulting correlation matrices were averaged across all eighteen slides. The spatial correlation matrix was clustered using hierarchical clustering to reveal genes that exhibit similar spatial expression patterns. We further assigned a color to each row and column in the matrix indicating the meta-module each gene belongs to.

### Code availability

Codes for data pre-processing, model training and evaluation were deposited into a public GitHub repository (https://github.com/gevaertlab/sequoia-pub).

### Data availability

Anonymized WSIs, gene expression and clinical data of the The Cancer Genome Atlas’ (TCGA) cohorts were retrieved from the publicly available Genomic Data Commons (GDC) portal (https://portal.gdc.cancer.gov). Gene expression data of the Clinical Proteomic Tumor Analysis Consortium (CPTAC) cohort were downloaded from GDC portal (https://portal.gdc.cancer.gov) and WSIs were obtained from the Cancer Image Archive with the accession URL (https://www.cancerimagingarchive.net/collections). Gene expression data and WSIs of the Tempus cohort were obtained through a data transfer agreement with Tempus Labs, Inc. Publicly available gene expression data and WSIs of The Genotype-Tissue Expression (GTEx) project were retrieved with the accession URL (https://gtexportal.org). The publicly available spatial transcriptomics data of GBM were acquired from Datadryad using the following accession URL (https://doi.org/10.5061/dryad.h70rxwdmj) [5]. The RNA-seq data and clinical annotations of the SCANB cohort were obtained from the accession URL (https://data.mendeley.com/datasets/yzxtxn4nmd/3), and data of the METABRIC cohort was obtained for cbioportal with accession URL (https://www.cbioportal.org/study/summary?id=brcametabric).

## Acknowledgments

Research reported here was further supported by the National Cancer Institute (NCI) under awards: R01 CA260271. The content is solely the responsibility of the authors and does not necessarily represent the official views of the National Institutes of Health. In addition, M. Pizurica was supported by a Fellowship of the Belgian American Educational Foundation and a grant from FWO 1161223N. F. Carrillo-Perez was also supported by a predoctoral scholarship from the Fulbright Spanish Commission. We are grateful for Roche Information Solutions (RIS) sponsorship, encouragement and support for this project.

## Appendix A Supplementary tables and figures

**Fig. A1:**
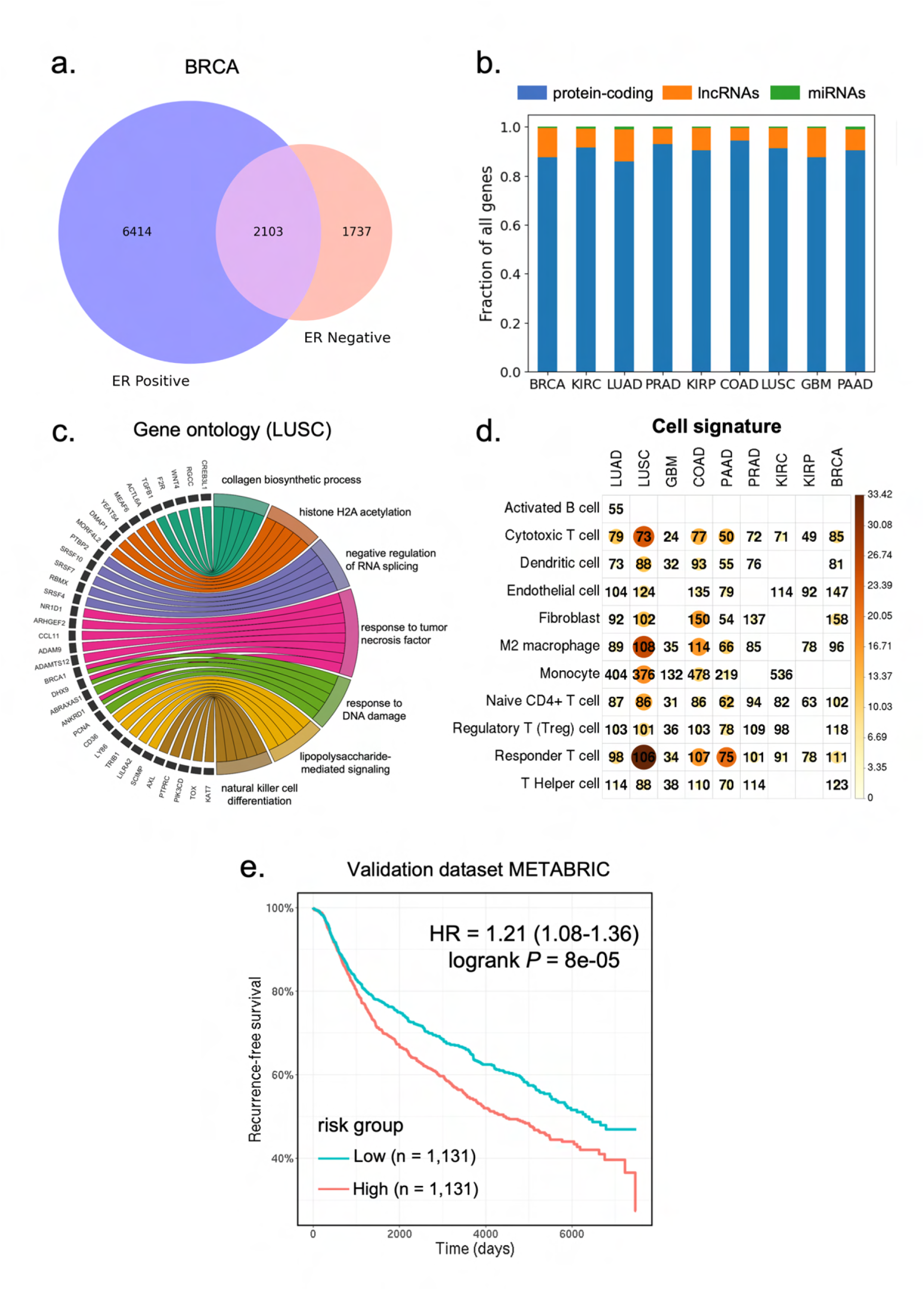
Characterization of the well-predicted genes. a) Venn diagram showing the number of well predicted genes in the estrogen-receptor (ER) positive and ER negative breast cancer. b) The proportion of protein-coding genes, miRNAs and lncRNAs among the well predicted genes from each cancer type. c) Circos plot showing the biological processes associated with the well predicted genes in LUSC. d) Heatmap showing the significant *P* values for the enrichment of cell-type signatures across cancer types. Color and size of the circles represent the negative log-transformed *P* values. Integers represent the absolute gene count in each category, and non-significant categories are left in blank. *P* values were adjusted for multiple testing using the Benjamini–Hochberg method. e) Kaplan-Meier curves of recurrence-free survival in the METABRIC validation dataset (n = 2,262 patients). Patients were split by the median risk score. HR: hazard ratio.

**Fig. A2:**
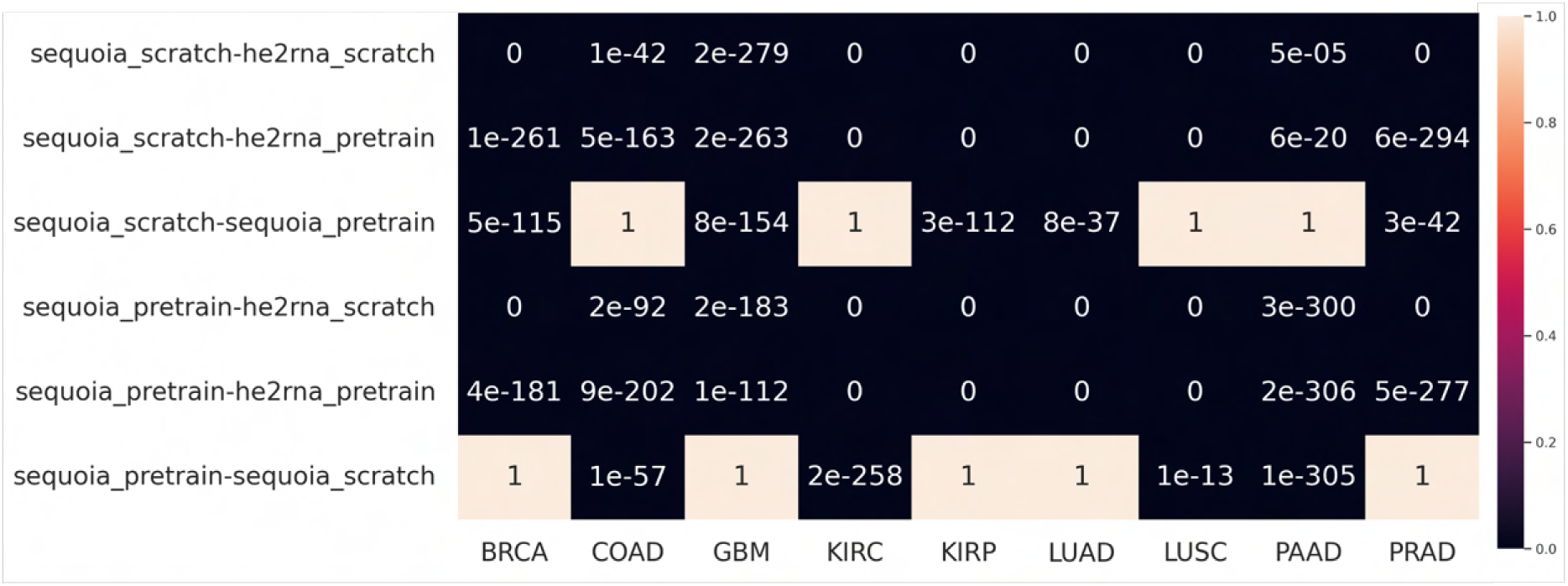
*P* values for testing the distributions of correlation coefficients for the top 1,000 most accurately predicted genes obtained from each model. A *P* value is calculated for each pairwise comparison using a one-sided Mann-Whitney U test for the hypothesis that model *x* is larger than model *y*, formatted on the left axis as *x*-*y*.

**Fig. A3:**
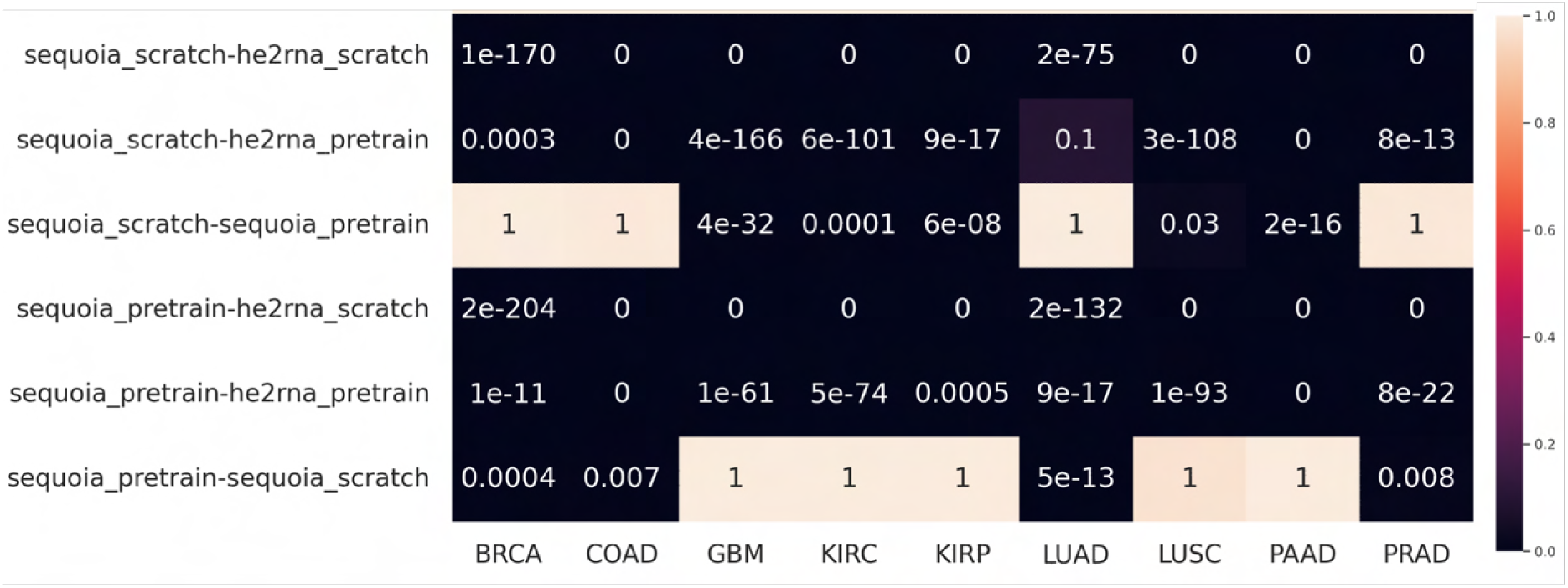
*P* values for testing the distributions of RMSE values between each of the two models. A *P* value was calculated for each pairwise comparison using a one-sided Mann-Whitney U test for the hypothesis that model *x* is smaller than model *y*, formatted on the left axis as *x*-*y*.

**Fig. A4:**
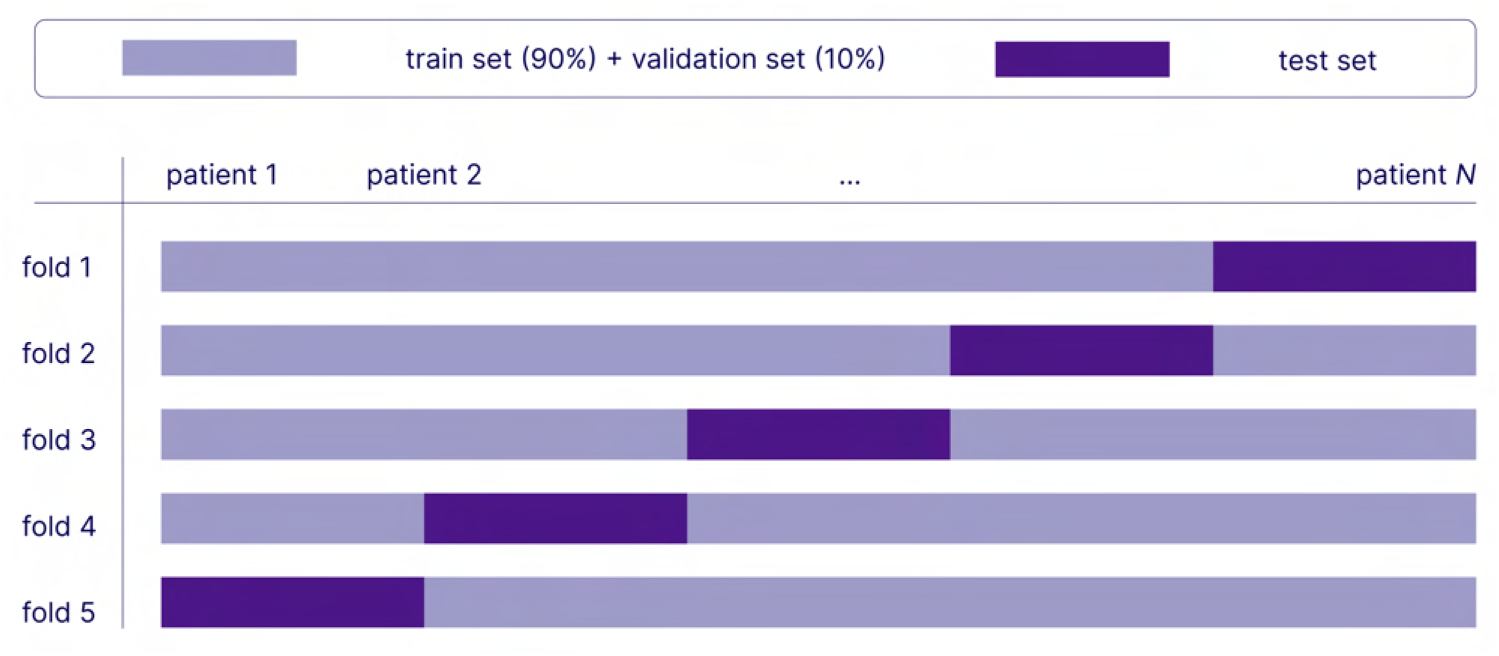
Data splitting. First, five folds are made each of which consist of a ‘global’ train and test set. The ‘global’ train set is further split into a train (90%) and validation (10%) set. In each fold *i*, validation set *i* is used to determine the optimal point to stop training model *i*, which is then evaluated on test set *i*. Afterwards, predictions on patients from test sets *i* (*i* = 1..5) are concatenated before calculating performance measures (e.g. Pearson correlation between predicted gene expression and ground truth expression across patients).

**Table A1:** Number of genes, Whole Slide Images and unique number of patients used from each cancer type in TCGA.

| TCGA project | number genes | number WSIs | number patients |
| --- | --- | --- | --- |
| TCGA-BRCA | 25,761 | 1,130 | 1,059 |
| TCGA-LUAD | 25,812 | 536 | 473 |
| TCGA-LUSC | 26,443 | 510 | 476 |
| TCGA-GBM | 26,530 | 237 | 102 |
| TCGA-PRAD | 25,587 | 448 | 401 |
| TCGA-PAAD | 26,172 | 202 | 176 |
| TCGA-KIRP | 25,141 | 299 | 275 |
| TCGA-KIRC | 26,653 | 514 | 508 |
| TCGA-COAD | 23,645 | 455 | 447 |
| total |  | 4,331 | 3,917 |

**Table A2:** Number of genes, Whole Slide Images and unique number of patients used from each cancer type in CPTAC.

| CPTAC project | number genes | number WSIs | number patients |
| --- | --- | --- | --- |
| CPTAC-BRCA | 25,761 | 106 | 106 |
| CPTAC-CCRCC | 26,653 | 302 | 211 |
| CPTAC-COAD | 23,645 | 103 | 103 |
| CPTAC-GBM | 26,530 | 94 | 94 |
| CPTAC-LUAD | 25,812 | 222 | 222 |
| CPTAC-LSCC | 26,443 | 109 | 108 |
| CPTAC-PDA | 26,172 | 146 | 146 |
| total |  | 1,081 | 989 |

**Table A3:**
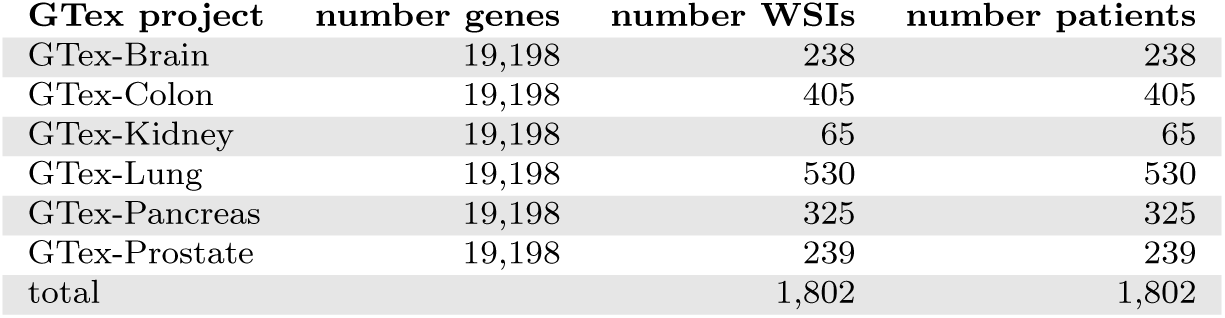
Number of genes, Whole Slide Images and unique number of patients used from each normal tissue in GTex.

**Table A4:** Number of genes significantly well predicted in the TCGA test sets. *N* shows the total number of slides available for each cancer type.

|  | he2rna_scratch | he2rna_pretrain | sequoia_scratch | sequoia_pretrain |
| --- | --- | --- | --- | --- |
| BRCA ($N = 1130$) | 1,429 | 2,902 | 12,239 | 11,069 |
| LUAD ($N = 536$) | 413 | 601 | 9,747 | 8,759 |
| KIRC ($N = 514$) | 0 | 66 | 6,985 | 10,086 |
| LUSC ($N = 510$) | 22 | 6 | 5,269 | 4,919 |
| COAD ($N = 455$) | 2,348 | 464 | 6,090 | 7,740 |
| PRAD ($N = 448$) | 973 | 2,124 | 8,895 | 10,825 |
| KIRP ($N = 299$) | 626 | 336 | 8,884 | 7,905 |
| GBM ($N = 237$) | 983 | 1,303 | 4,208 | 2,498 |
| PAAD ($N = 202$) | 314 | 387 | 418 | 3,378 |

**Table A5:** Median correlation coefficient between prediction and ground truth in TCGA test set for top 1000 genes within each model. Top genes defined as genes with highest correlation coefficient for each model type.

|  | he2rna_scratch | he2rna_pretrain | sequoia_scratch | sequoia_pretrain |
| --- | --- | --- | --- | --- |
| BRCA | 0.120 | 0.200 | 0.327 | 0.282 |
| LUAD | 0.116 | 0.117 | 0.284 | 0.269 |
| KIRC | 0.090 | 0.108 | 0.229 | 0.300 |
| LUSC | 0.097 | 0.088 | 0.199 | 0.206 |
| COAD | 0.223 | 0.141 | 0.268 | 0.310 |
| PRAD | 0.128 | 0.205 | 0.321 | 0.303 |
| KIRP | 0.164 | 0.135 | 0.368 | 0.333 |
| GBM | 0.188 | 0.205 | 0.289 | 0.242 |
| PAAD | 0.169 | 0.164 | 0.173 | 0.265 |

**Table A6:**
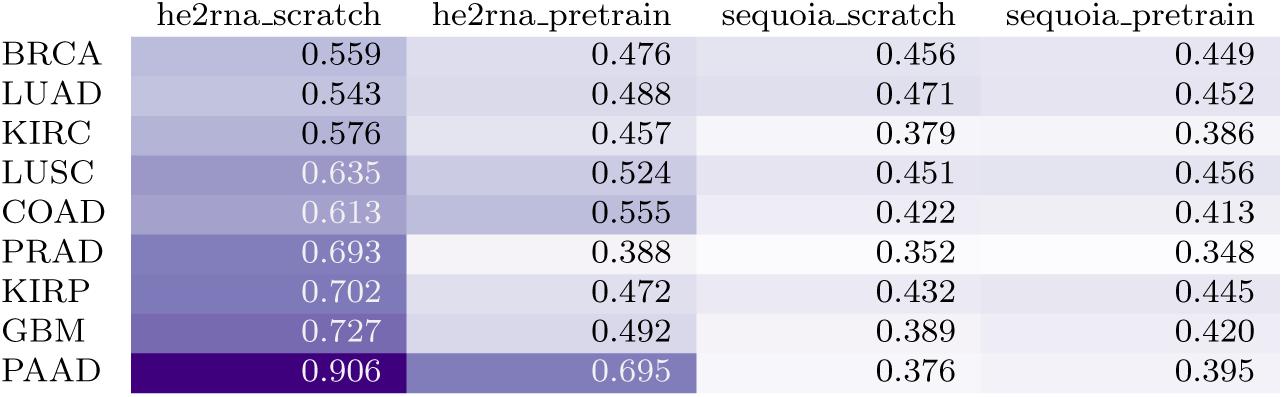
Median RMSE values between prediction and ground truth in TCGA test set within each model.

|  | he2rna_scratch | he2rna_pretrain | sequoia_scratch | sequoia_pretrain |
| --- | --- | --- | --- | --- |
| BRCA | 0.559 | 0.476 | 0.456 | 0.449 |
| LUAD | 0.543 | 0.488 | 0.471 | 0.452 |
| KIRC | 0.576 | 0.457 | 0.379 | 0.386 |
| LUSC | 0.635 | 0.524 | 0.451 | 0.456 |
| COAD | 0.613 | 0.555 | 0.422 | 0.413 |
| PRAD | 0.693 | 0.388 | 0.352 | 0.348 |
| KIRP | 0.702 | 0.472 | 0.432 | 0.445 |
| GBM | 0.727 | 0.492 | 0.389 | 0.420 |
| PAAD | 0.906 | 0.695 | 0.376 | 0.395 |

**Table A7:**
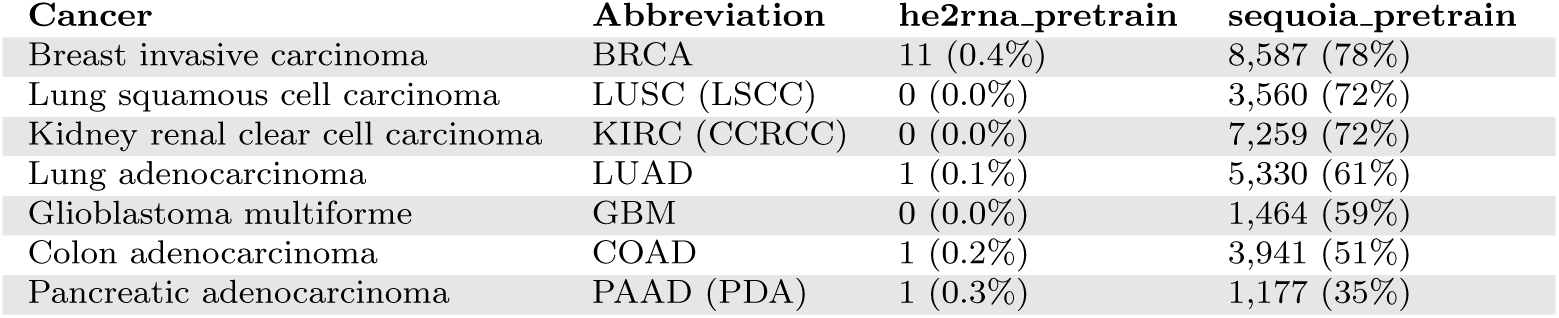
Number of genes validated in the CPTAC cohort using the *HE*2*RNA* model versus the *SEQUOIA* model.

**Table A8:** Median Earth Mover’s Distance between prediction and ground truth for top 500 genes from TCGA test set evaluated on different slides in spatial validation cohort.

| slide ID | median EMD | slide ID | median EMD |
| --- | --- | --- | --- |
| 242 | 0.116 | 266 | 0.096 |
| 243 | 0.152 | 268 | 0.238 |
| 248 | 0.170 | 269 | 0.128 |
| 251 | 0.171 | 270 | 0.134 |
| 255 | 0.113 | 275 | 0.132 |
| 259 | 0.123 | 296 | 0.138 |
| 260 | 0.131 | 304 | 0.121 |
| 262 | 0.126 | 313 | 0.179 |
| 265 | 0.174 | 334 | 0.197 |

## Notes

### Competing Interest Statement

The authors have declared no competing interest.

### Summary of Updates

Edits made to improve the abstract and introduction.

